# Measuring nonapoptotic caspase activity with a transgenic reporter in mice

**DOI:** 10.1101/196105

**Authors:** P. J. Nicholls, Thomas F. Pack, Nikhil M. Urs, Sunil Kumar, Yang Zhou, Gabriel Ichim, Joshua D. Ginzel, Gabor Turu, Evan Calabrese, Wendy L. Roberts, Ping Fan, Valeriy G. Ostapchenko, Monica S. Guzman Lenis, Flavio Beraldo, Jiri Hatina, Vania F. Prado, Marco A. M. Prado, Ivan Spasojevic, Joshua C. Snyder, Kafui Dzirasa, G. Allan Johnson, Marc G. Caron

## Abstract

The protease caspase-3 is a key mediator of apoptotic programmed cell death. But weak or transient caspase activity can contribute to neuronal differentiation, axonal pathfinding, and synaptic long-term depression. Despite the importance of sublethal, or nonapoptotic, caspase activity in neurodevelopment and neural plasticity, there has been no simple method for mapping and quantifying nonapoptotic caspase activity in rodent brains. We therefore generated a transgenic mouse expressing a highly sensitive and specific fluorescent reporter of caspase activity, with peak signal localized to the nucleus. As a proof of concept, we first obtained evidence that nonapoptotic caspase activity influences neurophysiology in an amygdalar circuit. Then focusing on the amygdala, we were able to quantify a sex-specific persistent elevation in caspase activity in females after restraint stress. This simple *in vivo* caspase activity reporter will facilitate systems-level studies of apoptotic and nonapoptotic phenomena in behavioral and pathological models.

**Significance Statement:** Caspase-3 is an enzyme that can cause cell death when highly active but can also perform important cellular functions, such as maturation and structural changes, when only weakly or transiently active. Despite the importance of this nonlethal type of caspase activity, there is no straightforward method to measure it in live rodents. We therefore developed mice that have a fluorescent reporter that is sensitive enough to detect nonlethal caspase activity. Surprisingly, we found that weak caspase activity influences the synchrony of neuronal firing across different brain regions. We also observed increased caspase activity in female mice after severe stress. This simple caspase activity reporter can subserve multiple applications in behavior and pathology research.

## Introduction

When caspase-3 is highly active, a proteolytic cascade ensues that typically ends in cell death by apoptosis (Yuan et al., 1993; Unsain & Barker, 2015). Weak or transient caspase-3 activity, however, is nonlethal and may be a key component of neurodevelopment by directing neuronal differentiation (Fernando et al., 2005), axonal pathfinding and branching (Westphal et al., 2010; Campbell & Okamoto, 2013), and synaptic metaplasticity (Chen et al., 2012). In the past several years mounting evidence indicates that, in the adult brain, nonapoptotic caspase activity (NACA) is an important contributor to neuronal plasticity, by driving NMDA receptor-mediated long-term depression (Li et al., 2010; Jiao & Li, 2011). Caspase activity in mature neurons may also underlie dendritic pruning (Ertürk et al., 2014). Uncovering the brain regions that display NACA at baseline and how NACA changes in response to perturbation will enable studies of its role in brain function.

Preclinical studies in cultured neurons (Du et al., 2004) and neonatal (Zhu et al., 2006; Renolleau et al., 2007), young adult, and aged rodents (Liu et al., 2009, 2011) have revealed sex-dependent differences in cell death pathways. In the context of hypoxia-ischemia, for example, cell death in females depends predominantly on caspase activity, while in males cell death occurs primarily by a caspase-independent mechanism (Zhu et al., 2006; Liu et al., 2009). This may be due to the more pronounced caspase-3 activation seen in females following hypoxia-ischemia and other cytotoxic insults (Liu et al., 2009; Du et al., 2004). Notably, these sex-dependent differences do not rely on estrogen levels or developmental stage (Liu et al., 2009). Although these studies were concerned with apoptotic caspase activity, they motivated us to investigate how sex influences baseline and stress-responsive NACA.

There is currently no tractable method for mapping and quantifying weak and transient caspase activity at cellular resolution throughout the brain. Such a method would enable systems-level studies of the relationship between NACA, neural circuits, and various behavioral and pathological models. Available transgenic fluorescent reporters of weak caspase activity that do not require reagents are limited to Drosophila (Bardet et al., 2008; Tang et al., 2015). In mice, the available caspase reporters rely on substrates for bioluminescence (Fu et al., 2013; Galbán et al., 2013) or beta-galactosidase activity (Khanna et al., 2010) which cannot be used for high-resolution studies of fixed tissues. And a FRET-based reporter in mice does not detect NACA (Yamaguchi et al., 2011). It was our goal to generate and validate a reagent-free caspase reporter in mouse brain that would produce a strong nuclear signal with minimal background activity.

Inteins are analogous to introns for proteins (Gogarten et al., 2002). In a process called protein splicing, conserved residues in the amino and carboxy terminals of an intein engage in thioester reactions that lead to the autocatalytic removal of the intein from flanking exteins. At the conclusion of protein splicing, the amino and carboxy terminal exteins are covalently bound to each other, linked by several residues from the ends of the removed intein. In a variant on this scheme, there are split inteins such that the amino and carboxy portions of a split intein recognize each other noncovalently, thereby facilitating the thioester reactions of the protein splicing process, again leading to covalently bound exteins as an end product (Shah & Muir, 2014). Inteins and split inteins are surprisingly widespread, having been identified in the genomes of roughly 50% of archaea and 25% of bacteria, as well as a few species of eukaryotes (Lennon & Belfort, 2017). Several bacterial split inteins have found application in synthetic biology, protein chemistry, and other fields. Of these, the split intein encoded by the dnaE gene of the cyanobacterium Synechocystis species PCC6803 has been well characterized and used in several contexts (Wu et al., 1998; Evans et al., 2000), including the reconstitution of split Cre recombinase in transgenic mouse brain (Wang et al., 2012). We chose this split intein when designing a caspase activity reporter based on a split version of the fluorophore mCerulean.

In this paper, we describe a novel transgenic reporter of caspase activity in mice. First, we demonstrate the simplicity of detecting apoptotic reporter signal in several models of programmed cell death. Then, to find an optimal brain region for studying NACA, we examine the relationship between caspase activity and neurophysiology in three brain areas. Finally, to demonstrate the utility of our reporter in measuring NACA *in vivo*, we focus on neurons in the anterior basolateral amygdala and find sex-dependent baseline and stress-induced differences in caspase activity. Taken together, our results suggest that NACA may underlie pathologic responses to severe behavioral stress.

## Materials and Methods

### CONTACT FOR REAGENT AND RESOURCE SHARING

Further information and requests for resources and reagents should be directed to and will be fulfilled by the Lead Contact, PJ Nicholls.

### EXPERIMENTAL MODEL AND SUBJECT DETAILS

#### Mice

All procedures involving animals were carried out in accordance with National Institutes of Health standards as approved by the Duke University Institutional Animal Care and Use Committee. The caspase activity reporter transgene (Figure 1A, annotated sequence available upon request) was synthesized and inserted into pCAGEN by GenScript. Pronuclear microinjection of the transgene into C57BL/6NHsd one-cell embryos was performed by Duke’s Transgenic Mouse Facility. Reporter mice used in experiments were male and female, primarily 8-10 weeks old but ranging from E17 to 4 months old, homozygous on a C57BL/6 (all except Figures 5D, 6, 7C, D) or mixed background (Figures 5D, 7C, D), and drug- and test-naïve. The mice in Figure 4E were P46 male caspase reporter hemizygous, floxed *Arpc3* homozygous, and PCP2-Cre wild-type or hemizygous. Finally, mice for the neurophysiology (Figure 6C-I) and pharmacokinetics (Figure 6A) experiments were 2-3-month-old male C57BL/6J from Jackson Labs. Mice were housed in a barrier facility on a 12/12 light cycle with species-specific heat and humidity and fed a standard laboratory diet with food and water *ad libitum*. Mice were group housed 2-5 per micro-isolator cage on corn cob bedding, and cages were changed every two weeks. For the *ex vivo* (Figure 3D, E), neurophysiology, and pharmacokinetics experiments, cagemates were allocated randomly to experimental groups. For the aBLA NACA quantification in Figure 7C, male and female cages were alternated in the order of perfusion; the first half of samples perfused were randomized separately from the second half of samples perfused; the randomization was double-blind; and the first half of samples were imaged in their new order followed by the second half in their new order. For the aBLA NACA quantification in Figure 7D, alternate mice within a cage were restrained; the order of perfusion alternated between control and restraint within cages and male and female between cages; and the randomization and order of imaging were as in Figure 7C.

**Figure 1.**
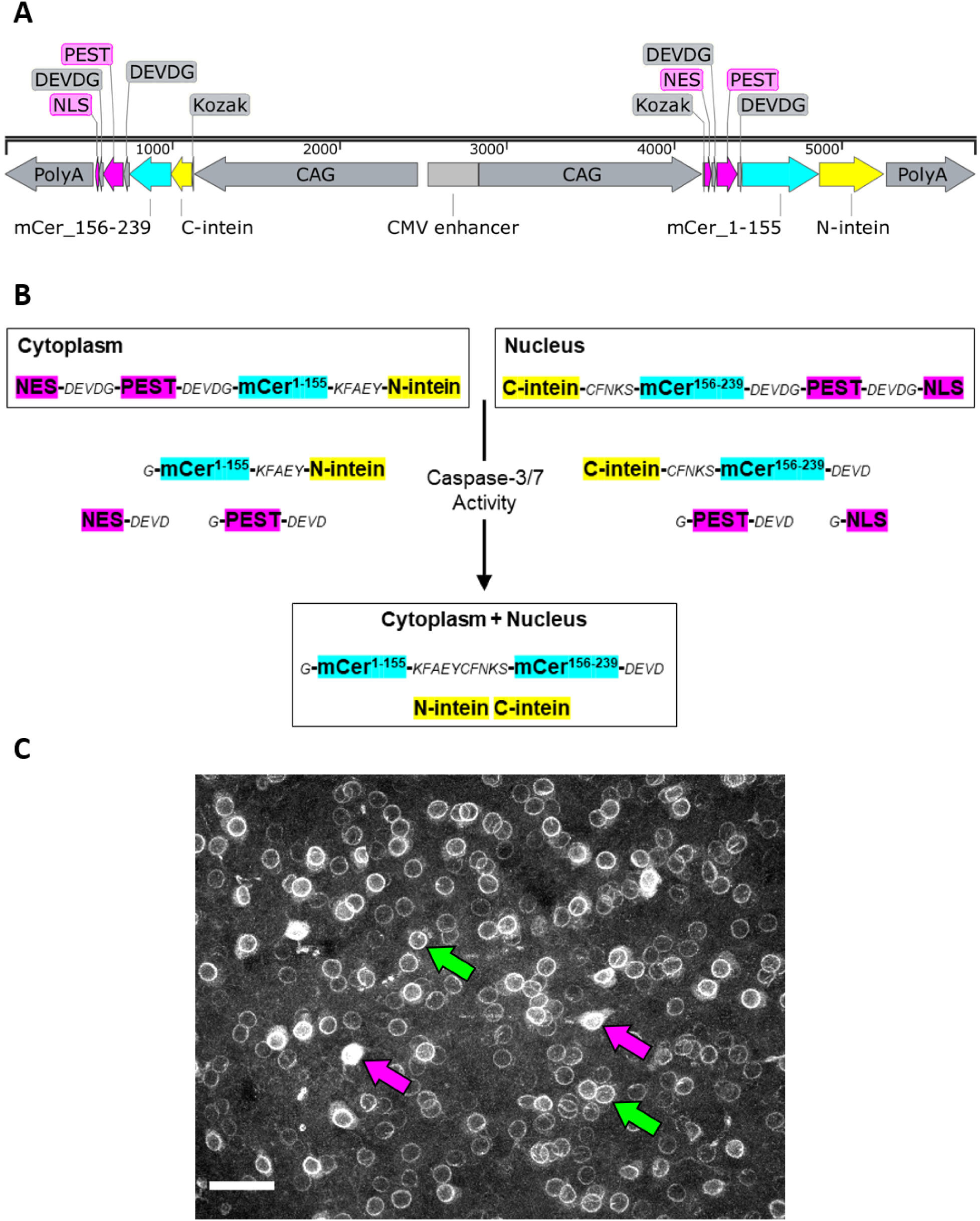
Design of a transgenic caspase-3/7 activity reporter. (A) Vector map. From left: PolyA, polyadenylation sequence; NLS, nuclear localization sequence; DEVDG, caspase cleavage site; PEST, protein degradation sequence; mCer, mCerulean fluorophore; C-, N-intein, carboxy- and amino-terminal bacterial inteins; Kozak, initiation sequence; CAG, CAGGS promoter; NES, nuclear export sequence. (B) Schematic of caspase- and intein-mediated bimolecular fluorescence complementation. KFAEY, CFNKS, intein linkers; other abbreviations as in (A). (C) Examples of weak reporter signal at nuclear envelope (green arrows) and brighter reporter signal filling nucleus and soma (magenta arrows). Basolateral amygdala; scale bar, 50 µm. See also Table 1.

**Figure 2.**
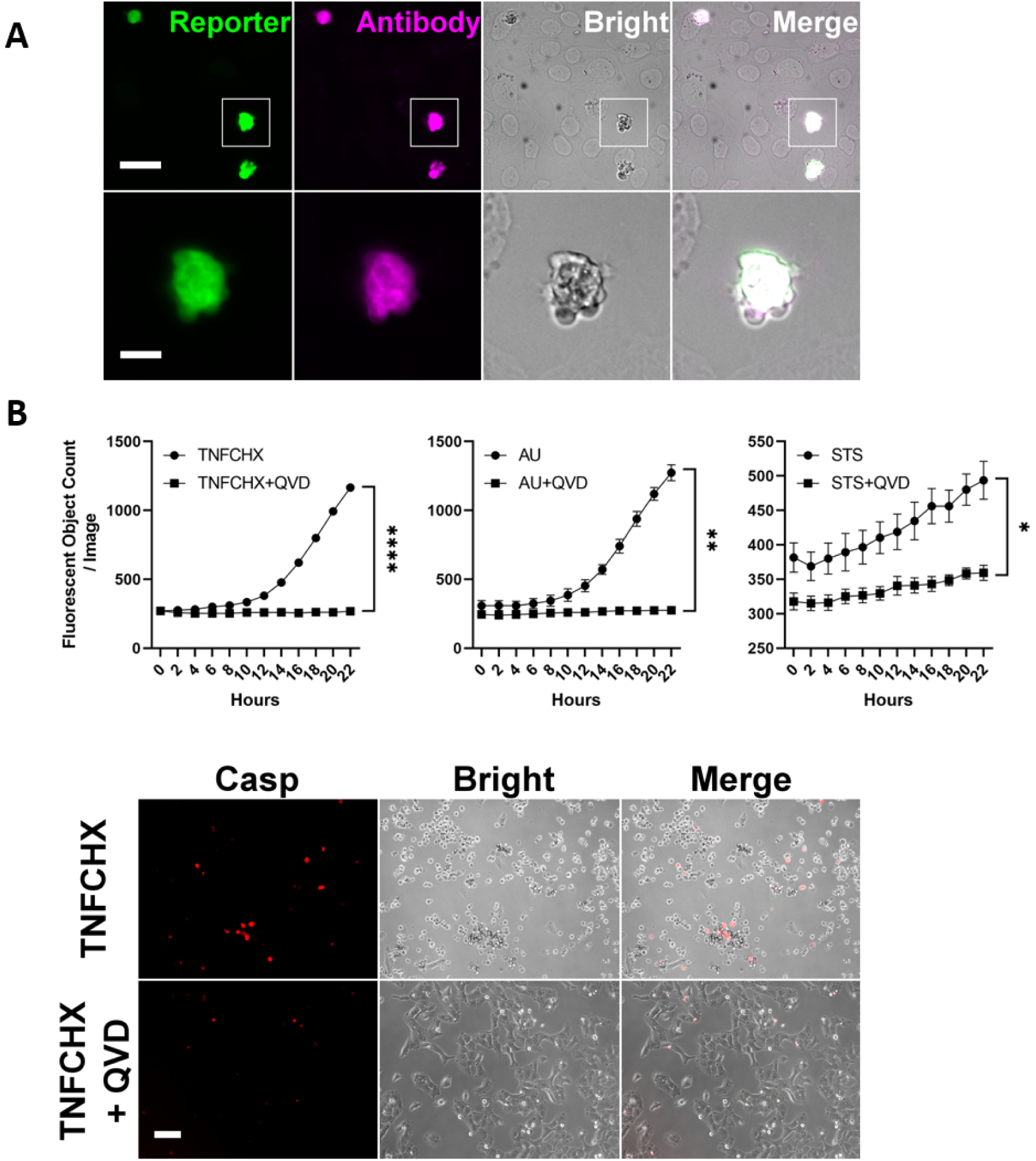
Detection of Apoptosis *In Vitro*. (A) In U2OS cells, transiently transfected caspase activity reporter (green) colocalized with cleaved caspase-3 antibody (magenta) in morphologically apoptotic cells (gray). Scale bar, 40 µm (above), 10 µm (below). (B) Above: A375 melanoma cells transfected with caspase activity reporter then treated with apoptotic stimuli showed an increase in fluorescent signal that was abolished by the caspase inhibitor Q-VD-OPh (QVD, 20 µM). TNFCHX, hTNFa 20 ng/ml + cycloheximide 1 µg/ml; AU, ABT-736 10 µM + UMI-77 10 µM; STS, staurosporine 1 µM. Two-way repeated-measures ANOVA (Treatment: left, F(1,4)=529.5; middle, F(1,4)=47.88; right, F(1,4)=10.78); mean ± SEM; n=3. *, p<0.05; **, p<0.01; ****, p<0.0001. Below: Example of *in vitro* conditions. Casp, caspase activity reporter. Scale bar, 100 µm. See also Table 1.

**Figure 3.**
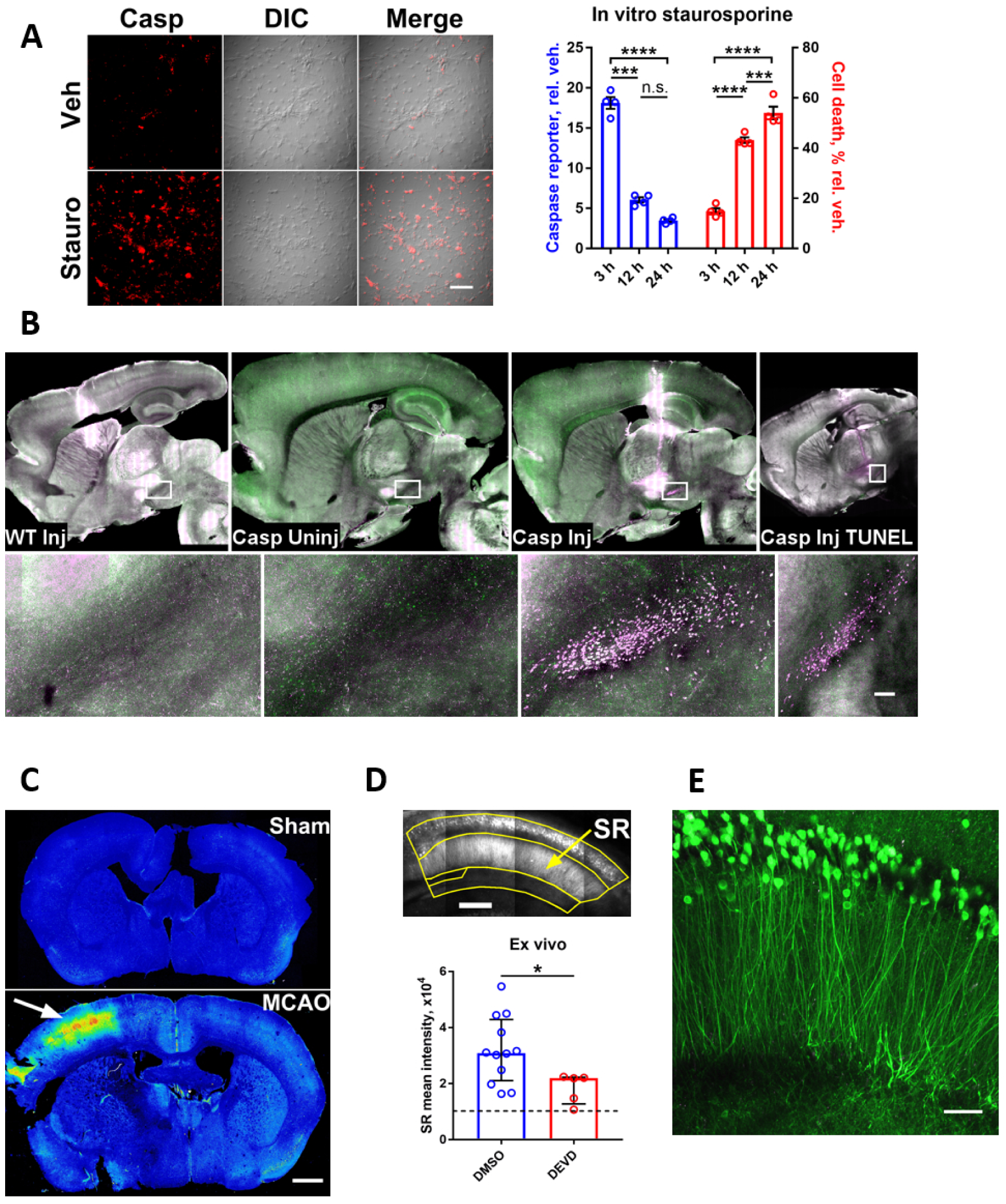
Validation of a transgenic caspase-3/7 activity reporter. (A) Representative data (left) and time course (right) of caspase reporter intensity and cell death in primary hippocampal neurons treated with 100-nM staurosporine. Two-way repeated-measures ANOVA with Tukey’s multiple comparisons test (Reporter x Time F(2, 12)=216.7); mean ± SEM; n=4. n.s., not significant; ***, p<0.001; ****, p<0.0001. Scale bar, 100 µm. (B) Apoptotic reporter signal (magenta) in the substantia nigra pars compacta ipsilateral to a 6-hydroxydopamine injection (Casp Inj), but not contralateral to the injection (Casp Uninj) or in injected wild-type (WT Inj), and confirmed by TUNEL stain (Casp Inj TUNEL). Green (Casp Uninj, Casp Inj) represents nonapoptotic reporter signal. Scale bar, 100 µm. (C) Caspase reporter distribution after middle cerebral artery occlusion (MCAO, arrow) versus sham. Red-blue is high-low signal intensity. Scale bar, 1 mm. (D) Attenuation of reporter signal by a caspase inhibitor (3.7-μM Z-DEVD-FMK) in acute hippocampal slices. SR, stratum radiatum. Dashed line denotes threshold of detection. Mann-Whitney test (U=9); median ± IQR; n=12 DMSO, 5 DEVD. *, p<0.05. Scale bar, 200 µm. (E) Hippocampal nonapoptotic reporter signal is predominantly in neurons of the stratum pyramidale. Acute hippocampal slice. Scale bar, 50 µm. See also Table 1.

**Figure 4.**
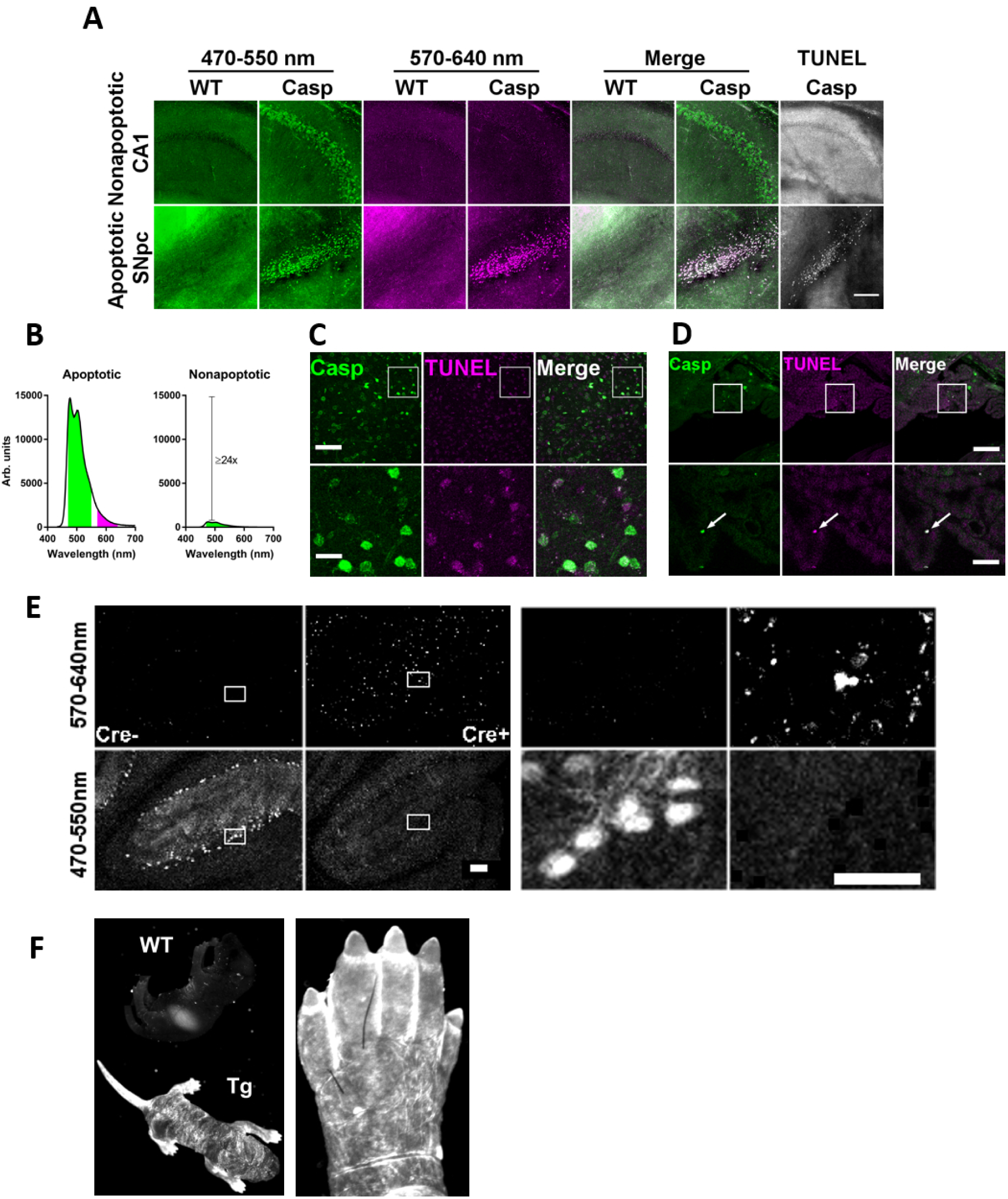
Separation of apoptotic and nonapoptotic reporter signal. (A) Separation of nonapoptotic (above) from apoptotic (below) reporter signal by comparing short- (green) and long-wavelength (magenta) emission. Data are the same experiment as Figure 3B. Scale bar, 200 µm. (B) Quantitative comparison of apoptotic and nonapoptotic reporter signal in short- (green) and long-wavelength (magenta) channels, derived from data used for Figure 7D (see Methods). (C) Lack of colocalization between nonapoptotic reporter signal (green) and TUNEL stain (magenta) in the basolateral amygdala. Scale bar, 100 µm (above), 25 µm (below). (D) Colocalization of reporter signal (green) with TUNEL stain (magenta) in apoptotic epithelial cells of the choroid plexus. Scale bar, 100 µm (above), 25 µm (below). (E) Separation of apoptotic (row 1) from nonapoptotic (row 2) reporter signal in Purkinje cells of *Arpc3* conditional knockout mice, with and without Cre recombination. Scale bar, 100 µm (left), 50 µm (right). (F) Caspase reporter signal is present in shedding skin of a transgenic P0 mouse (left, Tg) and in apoptotic webbing between its toes (right). Weak signal in wild-type (WT) P0 mouse is autofluorescence in the stomach. See also Table 1.

**Figure 5.**
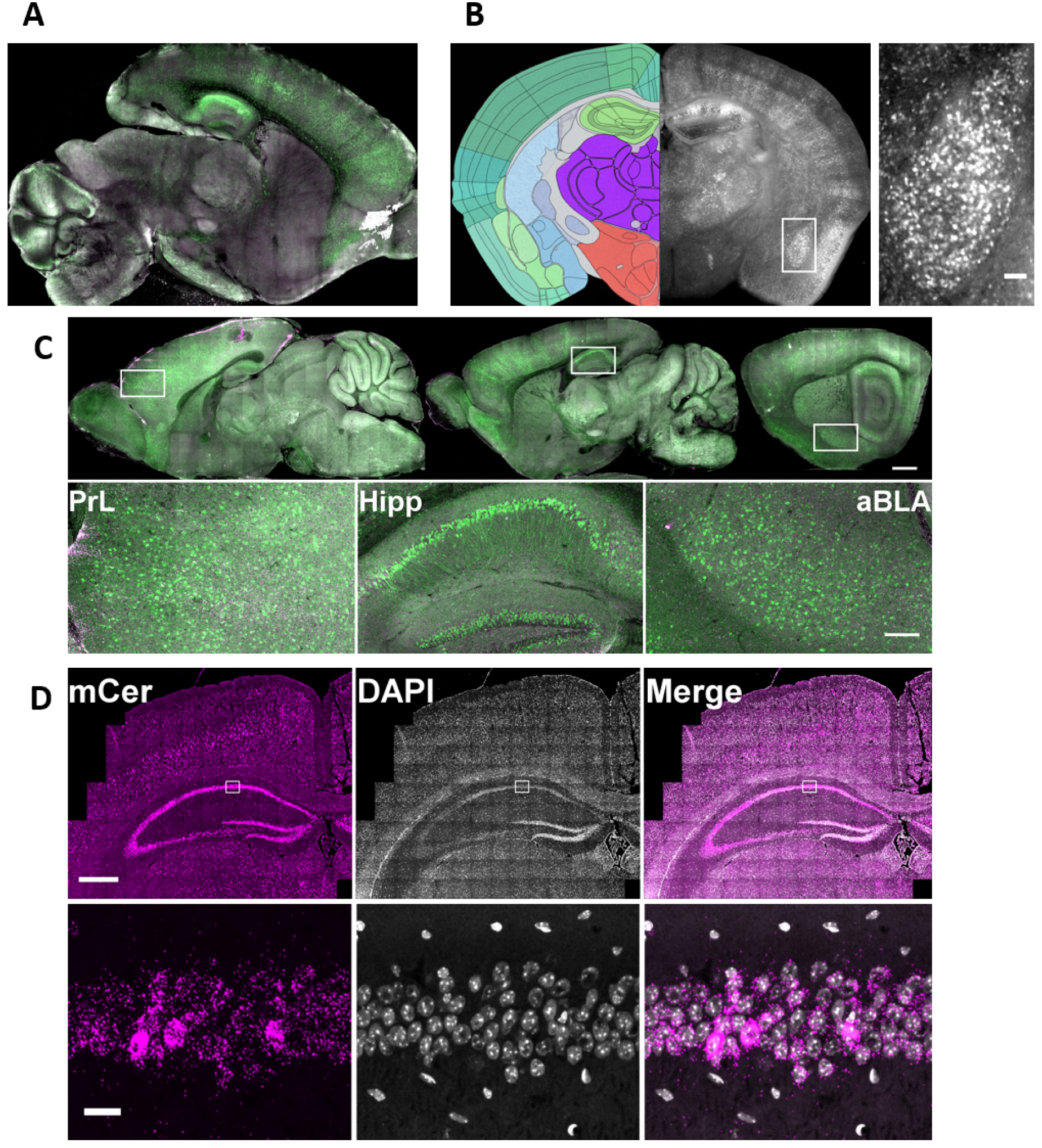
Regional and cellular location of nonapoptotic reporter signal. (A) Nonapoptotic reporter signal (green) in a clarified sagittal section of P22 brain. (B) Nonapoptotic reporter signal in a coronal section of P44 brain (right). Left atlas tracing image credit: Allen Institute (Lein et al., 2007). Inset: anterior basolateral amygdala. Scale bar, 100 µm. (C) Nonapoptotic signal (green) is elevated in prelimbic cortex (PrL), hippocampus (Hipp), and anterior basolateral amygdala (aBLA). Scale bar, 1 mm (above), 200 µm (below). (D) RNA *in situ* hybridization against mCerulean (mCer, magenta). In the insets, note lack of colocalization with DAPI signal outside of the pyramidal cell layer. Scale bar, 500 µm (above), 20 µm (below). See also Table 1.

**Figure 6.**
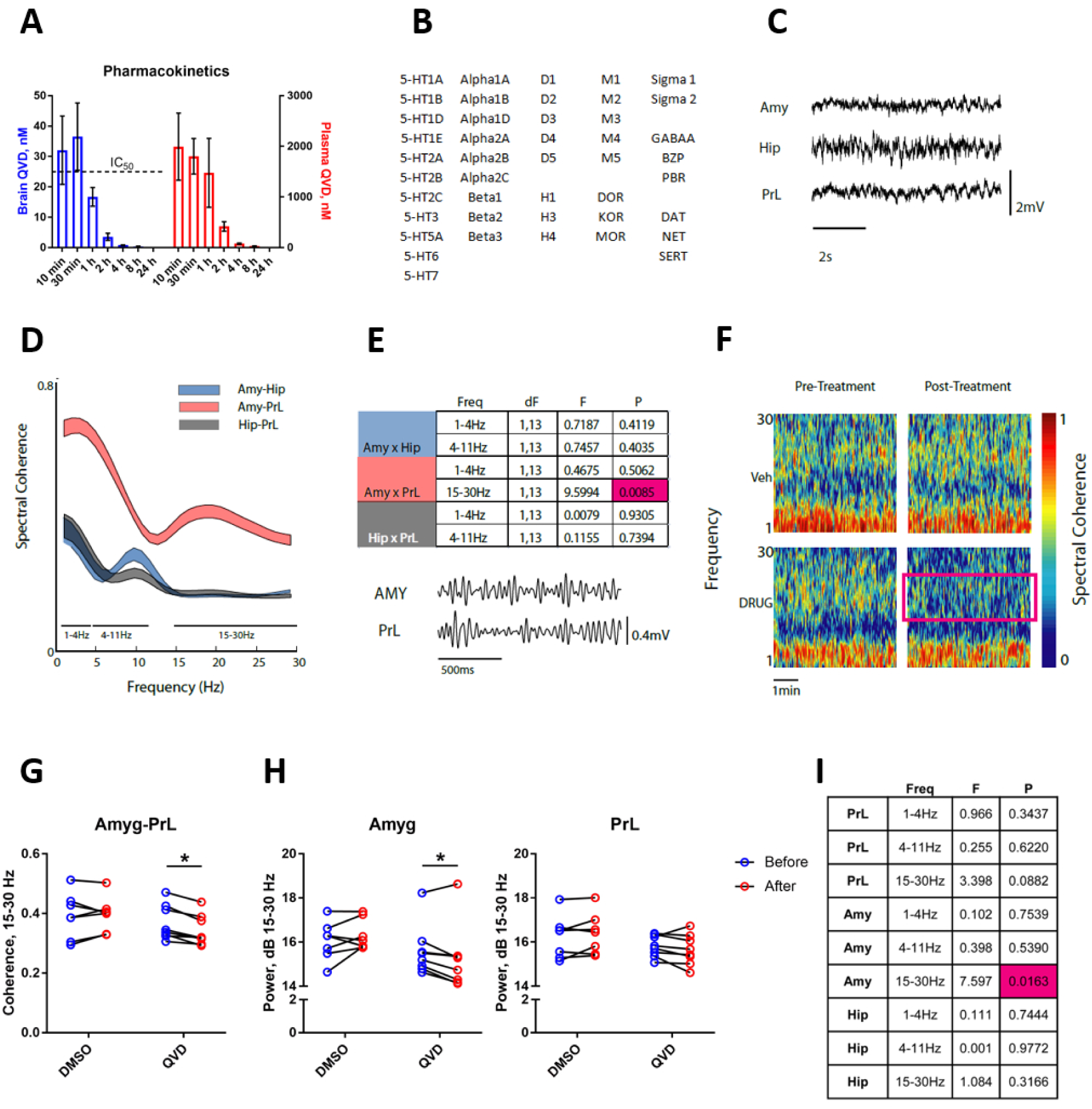
NACA contributes to functional connectivity in a stress-associated amygdalar circuit. (A) Pharmacokinetics of 5 mg/kg Q-VD-OPh in mouse brain and plasma after intraperitoneal injection. Mean ± SEM; 3-5 mice per time point. (B) Receptors and transporters screened for interaction with Q-VD-OPh. (C) Representative traces of local field potentials recorded from amygdala (Amy), hippocampus (Hip), and prelimbic cortex (PrL) in a freely behaving mouse. (D) Spectral coherence between three area pairs (Amy-Hip, Amy-PrL, and Hip-PrL). Data are presented as 95% confidence intervals; 15 mice. (E) Table of spectral coherence differences across functional connectivity bands depicted in (D), and representative β (15-30 Hz) frequency traces. Two-way repeated-measures ANOVA with Box-Cox transform; magenta highlights significant value. (F) Representative spectral coherence data between basal amygdala and prelimbic cortex. Magenta box shows decreased coherence after caspase inhibitor treatment. (G) Reduced functional connectivity between basal amygdala and prelimbic cortex after caspase inhibitor treatment (QVD). Two-way repeated-measures ANOVA with Sidak’s multiple comparisons test (Drug x Time F(1, 13)=6.594). (H) Decreased local field potential power after QVD treatment in basal amygdala (left), but not in prelimbic cortex (right). Two-way repeated-measures ANOVA with uncorrected Fisher’s LSD (Drug x Time: left, F(1, 13)=6.811; right, F(1, 13)=3.343). (I) Table of spectral power differences. Two-way repeated-measures ANOVA with Box-Cox transform; magenta highlights significant value. In (D, E, G-I), n=7 DMSO, 8 QVD; *, p<0.05.

**Figure 7.**
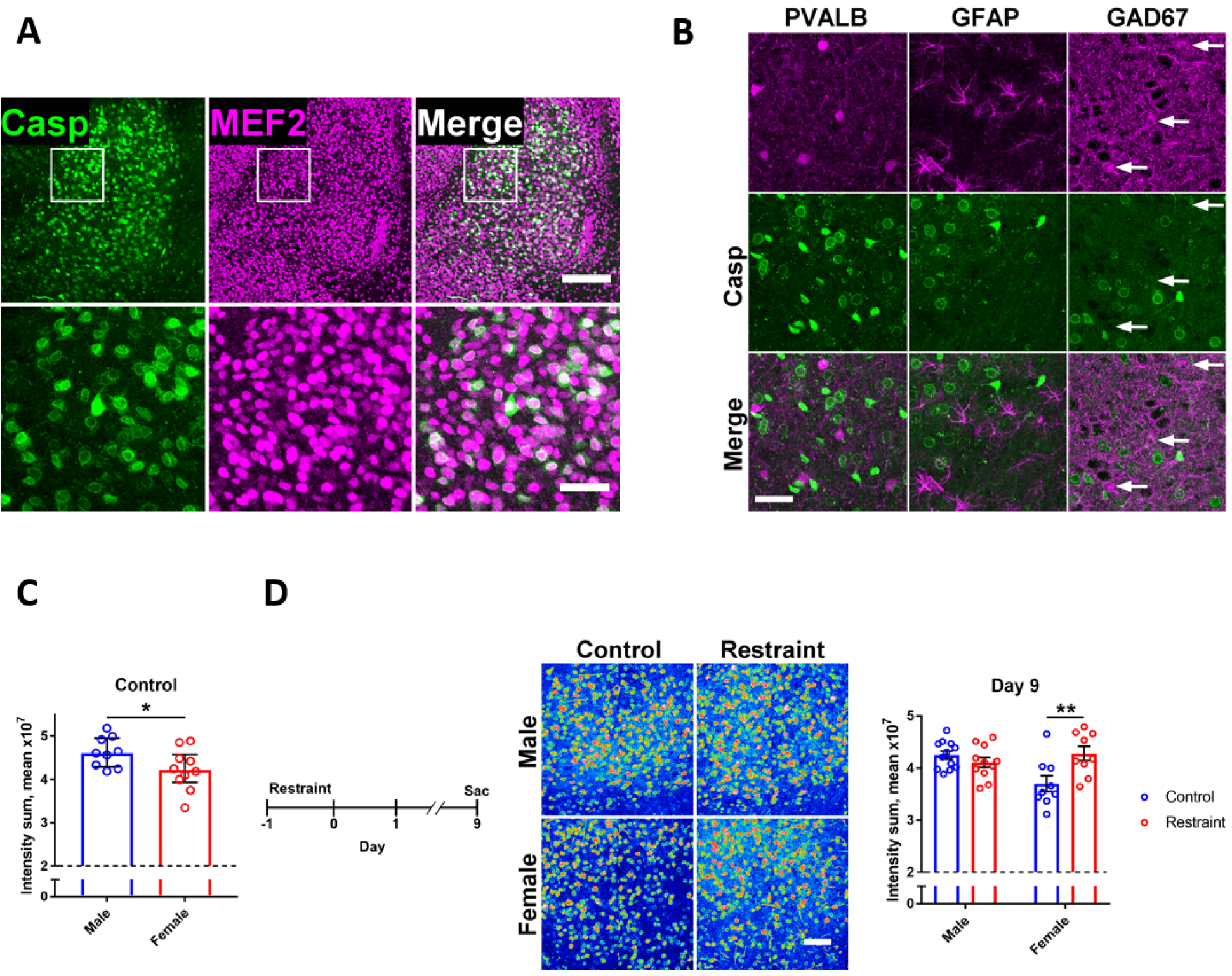
NACA, the amygdala, and behavioral stress. (A) Colocalization of reporter signal (Casp, green) with the neuronal marker myocyte enhancer factor 2 (MEF2, magenta) in basolateral amygdala. Scale bar, 200 µm (above), 50 µm (below). (B) Lack of colocalization of reporter signal (Casp, green) with (all magenta): parvalbumin (PVALB), glial fibrillary acidic protein (GFAP), and glutamic acid decarboxylase 67 kDa (GAD67). White arrows indicate cells with elevated antibody staining. Scale bar, 50 µm. (C) Reduced baseline NACA in female mice. Dashed line represents threshold of detection. Mann-Whitney test (U=20); median ± IQR; n=9 M, 10 F. (D) Left: Time course of the experiment. Middle: Representative caspase reporter intensity data (red-blue, high-low intensity). Right: Sex-specific increase in NACA 9 days after restraint stress. Dashed line denotes threshold of detection. Two-way ANOVA with Sidak’s multiple comparisons test (Sex x Stress F(1, 37)=9.931); mean ± SEM; n=12 M Cont., 11 M Rest., 9 F Cont., 9 F Rest. from 2 independent experiments. Scale bar, 100 µm. *, p<0.05; **, p<0.01. See also Table 1.

**Table 1.** Imaging parameters. Related to Figures 1–5 and 7.

| Figure | Specimen | Pixel Size (μm) | Z Interval (μm) | Slices | Projection | Channels | Excitation (nm) | Emission (nm) | System |
| --- | --- | --- | --- | --- | --- | --- | --- | --- | --- |
| 1C | Fixed | 0.41 | 2 | 76 | Maximum | Subtracted | 858 | 470-550, 570-640 | Ti:sapphire laser, 20x/1.0NA |
| 2A | Fixed | 0.18 | 0 | 1 | N/A | Single, merged | 424-448 (green), 538-562 (magenta), LED (gray) | 460-500 (green), 570-640 (magenta), transmitted light (gray) | Mercury arc lamp and LED, 2.3x/0.57NA |
| 2B | In vitro | 0.64 | 0 | 1 | N/A | Single, merged | 450-490 (red), LED (gray) | 500-550 (red), transmitted light (gray) | Mercury arc lamp and LED, 10x/0.3NA |
| 3A | In vitro | 0.58 | 0 | 1 | N/A | Single, merged | 405 (red), 488 (gray) | 430-470 (red), transmitted light (gray) | Diode laser (red), Ar laser (gray), 10x/0.4NA |
| 3B Left 3 | Clarified | 3.29 | 20 | 51 | Maximum | Merged | 858 | 470-550, 570-640 | Ti:sapphire laser, 20x/1.0NA |
| 3B TUNEL | Declarified | 3.29 | 20 | 11 | Maximum | Merged | 1000 | 470-550, 570-640 | Ti:sapphire laser, 20x/1.0NA |
| 3C | Fixed | 2.49 | 0 | 1 | N/A | Single | 458 | 480-495 | Ar laser, 10x/0.4NA |
| 3D | Acute slice | 0.82 | 10 | 15 | Sum | Subtracted | 858 | 470-550, 570-640 | Ti:sapphire laser, 20x/1.0NA |
| 3E | Acute slice | 0.2 | 2 | 21 | Maximum | Merged | 858 | 470-550, 570-640 | Ti:sapphire laser, 20x/1.0NA |
| 4A left 6 | Clarified | 3.29 | 20 | 51 | Maximum | Single, merged | 858 | 470-550, 570-640 | Ti:sapphire laser, 20x/1.0NA |
| 4A TUNEL | Declarified | 3.29 | 20 | 11 | Maximum | Merged | 1000 | 470-550, 570-640 | Ti:sapphire laser, 20x/1.0NA |
| 4C | Fixed | 0.82 | 5 | 41 | Maximum | Single, merged | 858 | 470-550, 570-640 | Ti:sapphire laser, 20x/1.0NA |
| 4D | Fixed | 0.82 | 5 | 41 | Maximum | Single, merged | 858 | 470-550, 570-640 | Ti:sapphire laser, 20x/1.0NA |
| 4E | Fixed | 0.82 | 10 | 21 | Maximum | Single (row 1), subtracted (row 2) | 858 | 470-550, 570-640 | Ti:sapphire laser, 20x/1.0NA |
| 4F left | In vivo | 9.29 | 0 | 1 | N/A | Single | 424-448 | 460-500 | Mercury arc lamp, 1x/0.25NA |
| 4F right | In vivo | 9.29 | 0 | 1 | N/A | Single | 424-448 | 460-500 | Mercury arc lamp, 1x/0.25NA |
| 5A | Clarified | 3.29 | 20 | 51 | Maximum | Merged | 858 | 460-500, 520-550 | Ti:sapphire laser, 20x/1.0NA |
| 5B | Fixed | 2.27 | 0 | 1 | N/A | Single | 450-490 | 500-550 | Mercury arc lamp, 2.3x/0.57NA |
| 5C | Fixed | 3.29 | 20 | 14 | Maximum | Merged | 858 | 470-550, 570-640 | Ti:sapphire laser, 20x/1.0NA |
| 5D | Fixed | 0.26 | 2 | 12 | Maximum | Single, merged | 405 (gray), 561 (magenta) | 415-470 (gray), 570-607 (magenta) | Diode (gray), DPSS (magenta), 40x/1.2NA |
| 7A | Fixed | 0.82 | 5 | 51 | Maximum | Single, merged | 858 | 470-550 (green), 570-640 (magenta) | Ti:sapphire laser, 20x/1.0NA |
| 7B | Fixed | 0.82 | 5 | 17 (col 1), 12 (col 2), 3 (col 3) | Maximum | Single, merged | 858 | 470-550 (green), 570-640 (magenta) | Ti:sapphire laser, 20x/1.0NA |
| 7D | Fixed | 0.55 | 5 | 51 | Maximum | Subtracted | 858 | 470-550, 570-640 | Ti:sapphire laser, 20x/1.0NA |

#### Primary Cultures (Figure 3A)

Primary cultures of hippocampal neurons (Figure 3A) were obtained from caspase reporter pregnant females on day E17 as previously described (Beraldo et al., 2010, 2013). Briefly, mouse hippocampal neurons of both sexes were harvested at day 17 of embryonic development (E17) and maintained in dishes coated with poly-L-lysine (Sigma-Aldrich, Canada) for 7 days in Neurobasal medium (Invitrogen) with 2% B-27 supplement (Invitrogen). Half of the medium was replaced every 2-3 days. All procedures were conducted in accordance with approved animal use protocols at the University of Western Ontario (2016/103) following Canadian Council of Animal Care and National Institutes of Health guidelines.

### METHOD DETAILS

#### Real-time PCR

Quantitative PCR for genotyping was performed with SYBR Select Master Mix (Life Technologies), 400-nM primers (synthesized by Integrated DNA Technologies), and a LightCycler (Roche). Later genotyping was performed on a StepOnePlus system (Applied Biosystems) with PowerUp SYBR Green Master Mix (Thermo Fisher Scientific #A25742) and 500-nM primers (IDT). Genotyping primers: CMV enhancer forward, CCCACTTGGCAGTACATCAA; CMV enhancer reverse, GCCAAGTAGGAAAGTCCCATAA. Control primers: Actb intron forward, GCCGTATTAGGTCCATCTTGAG; Actb intron reverse, GGTTTGGACAAAGACCCAGA.

#### Microscopy

We used a Zeiss Axio Zoom.V16 for Figures 2A, 4F, 5B; an Incucyte Zoom system (Sartorius) for Figure 2B; an Olympus confocal FV1000 for Figure 3A, C; a Zeiss LSM 880 for Figure 5D; and a Zeiss LSM 510 with an automated stage, Coherent Chameleon laser, and halogen lamp for all other microscopy. In two-color images, green represents shorter wavelengths and magenta longer wavelengths, as detailed in Table 1. Images were processed in ImageJ. For subtracted images, the magenta channel was linearly subtracted from the green channel, to remove autofluorescence and apoptotic reporter signal. Only linear window-level adjustments were made to images; and images within a figure panel were adjusted as a single image, except for the TUNEL stain in Figures 3B, 4A. In Figures 3B, 4E, a linear unbiased MATLAB script (available upon request) was used to correct for a stable shading abnormality in our multiphoton system. Clarification was performed as previously described (Hama et al., 2011).

#### Immunocytochemistry (Figure 2A)

U2OS cells were seeded onto 35 x 10 mm tissue culture dishes (Genesee) pretreated with poly-D-lysine at a density of 5 × 10^5^ cells. The next day, cells were transfected with 2.5 µg of Apo plasmid using Lipofectamine 2000 (Thermo Fisher). The media was replaced with 2 ml Opti-MEM (Thermo Fisher) 5 hours after transfection. After 40 hours, cells were washed with PBS, fixed for 10 minutes in formalin at room temperature, and incubated overnight at 4 C in 1:500 cleaved caspase-3 antibody (Cell Signaling Technology #9661). The following day, cells were washed with PBS and incubated for 2 hours at room temperature in 1:500 Alexa Fluor 568 (Thermo Fisher #A11011). After final washes in PBS, the cells were immediately imaged on an Axio Zoom.V16 (Zeiss).

#### Live Cell Imaging (Figure 2B)

A375 melanoma cells were seeded onto 10 cm dishes at a density of 1.5 x 10^6^, transfected 24 hours later with 10 µg of Apo plasmid using Lipofectamine 2000, and another 24 hours later reseeded into 48 well plates at 5 x 10^4^ cells per well. After another 24 hours, cells were treated with apoptotic stimuli; then plates were imaged every 2 hours using an Incucyte Zoom system (Sartorius) to quantify the number of fluorescent cells per field of view.

Separately, for the images in Figure 2B, A375 melanoma cells were transiently transfected with the Apo plasmid, then plated 24 hours later in coated dishes (Ibidi), and then treated another 24 hours later with hTNFa (20 ng/ml) + cycloheximide (1 µg/ml) alone or with the pan-caspase inhibitor Q-VD-OPh (20 µM) for 24 hours. Images were captured by epifluorescence microscopy and transillumination.

#### Hippocampal Cultures (Figure 3A)

On DIV 7, hippocampal neurons were treated with either vehicle or staurosporine (100 nM). Cells were imaged at 3, 12 and 24 hours after treatment, and caspase-3 activity was measured using fluorescence from the mCerulean reporter. Fluorescence of neuronal cultures from 4 independent embryos of either sex was quantified using an FV1000 Olympus confocal microscope (excitation 405 nm and emission 470-485 nm). Integrated fluorescence was divided by total cell number and normalized to control cells that were not subjected to staurosporine, using ImageJ software (NIH, Bethesda, MD).

#### 6-Hydroxydopamine Microinjections (Figures 3B, 4A)

In mice under constant isoflurane anesthesia, 1 μl of 6-hydroxydopamine (3 mg/ml) dissolved in saline containing 0.02% ascorbic acid was injected at coordinates AP=−1.2, ML=−1.2, and DV=−4.75 with a Hamilton syringe fitted with a pulled glass capillary needle at 0.4 μl/min using a Nanojet syringe pump. Fourteen hours after injection, mice were transcardially perfused with 4% PFA, and brains were fixed overnight in 4% PFA at 4 C. The ApopTag TUNEL stain kit (Millipore #S7165) was used for Figures 3B, 4A.

#### Middle Cerebral Artery Occlusion (Figure 3C)

Three- to 4-month-old male caspase reporter homozygotes were subjected to 2-hour middle cerebral artery occlusion (MCAO) or were sham-operated. After 24-hour recovery, mice were perfused with 4% PFA in PBS. Perfused brains were incubated overnight at 4 C, after which 50-µm-thick coronal sections were prepared using a vibratome. Slices were immediately mounted with PBS on glass slides and imaged using an Olympus confocal FV1000 equipped with a 10x/0.4NA objective.

#### Acute Hippocampal Specimens (Figure 3D, E)

For *ex vivo* specimens, 3-4-week-old mice were anesthetized at time 0 and transcardially perfused with chilled oxygenated artificial CSF (in mM: NaCl 119, KCl 2.5, NaHCO_3_ 26, NaH_2_PO_4_ 1.25, glucose 11, CaCl_2_ 0.5, MgSO_4_ 1.3); then a vibratome-cut hemisphere was maintained in oxygenated artificial CSF with CaCl_2_ 2 mM during multiphoton scanning. Z-DEVD-FMK was purchased from Millipore (#264156).

#### Immunohistochemistry (Figures 3B, 4A, C, D, 7A, B)

Tissue stains were performed with 1:500 primary antibody and 1:500 Alexa Fluor 594 secondary antibody (Thermo Fisher), both overnight at 4 C. Primary antibodies were: MEF2D (Millipore #AB2263), Parvalbumin (Swant #PVG213), GFAP (Cell Signaling #3670), GAD67 (Millipore #MAB5406). The ApopTag TUNEL stain kit (Millipore) was used for Figures 3B, 4A, C, D.

#### Apoptotic vs. Nonapoptotic Signal Intensity (Figure 4B)

Based on the Cerulean emission spectrum available at http://tsienlab.ucsd.edu/Documents.htm and our observation that the 16-bit intensity mean of apoptotic reporter signal is 65535 in the 570-640-nm channel (0.06133 of total spectrum), and our calculation of 34555.6 for the mean 16-bit intensity mean in the 470-550-nm channel (0.7745 of total spectrum) for nonapoptotic signal in the 23 samples of one replicate of the experiment in Figure 7D, we found (65535/0.06133)/(34555.6/0.7745) = 23.9

And since this value is limited by the maximal 16-bit intensity mean of 65535, the true quotient of apoptotic to nonapoptotic signal intensity would be greater than or equal to 24.

#### mRNA Fluorescence *In Situ* Hybridization (Figure 5D)

Advanced Cell Diagnostics RNAscope® Multiplex Fluorescent Kit (323100) was used to detect mCerulean mRNA expression in fixed frozen sections. Sections were prepared and pretreated before hybridization and signal development of the mCerulean probe (ACD Bio. 1108841-C1) following the Multiplex Fluorescent Kit user manual. Probe labeling was done with Opal 570 dye (PerkinElmer, Inc NEL801001KT). Tiled images were acquired using a Zeiss LSM 880 with 10X and 40X objectives.

#### Measuring aBLA NACA (Figure 7C, D)

For the stress intervention (Figure 7D), ∼8-week-old male and female caspase reporter homozygotes housed 2-5 per cage were transferred to a nearby experimental procedure room within their barrier facility, where they acclimated overnight. At approximately 07:00 the next day, half of the mice were randomly selected and inserted into 50-ml tubes that had 0.25-in holes for breathing. For cages with an odd number of mice, selection was designed so that the total number of mice in each of the 4 experimental groups would be nearly equivalent. Immediately after a mouse was placed in a tube, it was returned to its cage where it stayed with its restrained and unrestrained cagemates. All mice were deprived of food and water for the next 24 hours. Mice were observed, and tubes were rolled periodically to assess responsiveness. No mice became unresponsive during the test. Twenty-four hours later, restrained mice were released from their tubes, washed briefly in a room-temperature water bath, and returned to their cages with their unrestrained cagemates. For the next 9 days, all mice had access to food and water *ad libitum*. The released mice were again observed for distress or unresponsiveness, which were not detected during this test. Within a few hours of being released, restrained mice were indistinguishable from unrestrained mice upon gross inspection. After 9 days, all mice were transferred to a separate facility for transcardial perfusion.

Mice were perfused in order, such that restrained mice alternated with unrestrained mice and male cages alternated with female cages. Immediately after being weighed, each mouse was administered a mixture of ketamine 100 mg/kg, xylazine 10 mg/kg, and heparin 5000 U/kg in saline and given 10-15 minutes to attain a surgical plane of anesthesia. Mice were observed and tested by toe pinch for inadequate anesthesia, which was not noted in this experiment. No mice were given an additional dose of anesthetic. After 10-15 minutes of anesthesia, each mouse was pinned to a board in a chemical fume hood and rapidly dissected to reveal the liver and heart. Then the apex of the left ventricle was pierced with a 25-gauge needle attached to a 25-ml syringe containing 25 ml of 1:1 10% neutral buffered formalin : PBS plus 100 μl of 1000 U/ml heparin. Immediately, the solution was forced into the circulation at a moderate pace, and the right atrium was snipped as it expanded. All 25 ml of formalin/PBS/heparin was forced through the circulation at a steady pace within 1-2 minutes. Each mouse was assessed for paleness of liver and stiffness of body; 1 mouse out of the 42 used in the stress experiment was determined to have a poor perfusion based on these criteria and was later excluded from data analysis before the randomization was unblinded.

Immediately after perfusion, the skin and mandible were carefully dissected from the skull, such that the skull was not compressed or dropped during the procedure. The skull was carefully placed in a 50-ml tube containing 25 ml of the perfusate solution, and each tube was placed on its side in a box on a rotator at ∼60 rpm in a 4 C cold room for 48 hours. During this time, numbers on the tubes were replaced by letters in a random order by a person blind to the numbering system. The letter assignments of the numbers were known only by this individual, and the information was shared (i.e., the double-blind was broken) only after the completion of data analysis. After the tubes had been in the cold room for 2 days, brains were carefully extracted from skulls and placed in PBS. Brains were then cut into 200-μm coronal sections in PBS on a vibratome under dim room light. Sections corresponding to the aBLA were transferred to individual wells in a 24-well plate containing 1 ml of PBS/1% Triton X-100 per well. Plates were stored overnight in the dark in a 4 C cold room. The next day, sections were examined in plates by brief halogen transillumination on a dissecting microscope, and the ideal section for aBLA imaging was determined for each mouse in a blinded manner. Plates were then returned to the dark in a 4 C cold room for overnight storage.

The next day, 3 sections were mounted per gelatin-coated slide, with 150-μm square coverslips Scotch Magic-taped to both ends supporting a 170-μm coverslip again secured by a single layer of Scotch Magic tape. Sections were kept wet with PBS/1% Triton X-100. Each slide was Krazy-glued to the bottom of a 10-cm tissue culture dish which was then filled with 20 ml of PBS/1% Triton X-100 and placed in the dark at room temperature. Samples were transported to a multiphoton microscope, and an additional 20 ml of PBS/1% Triton X-100 was added to each plate. Then a single 2×2 tiled image centered on the predetermined ideal aBLA was obtained for each specimen according to the parameters listed in Table 1. Multiphoton scans were analyzed with Imaris (Bitplane) to quantify the reporter signal in each nucleus using the Spots function with the following parameters: Estimated Diameter = 15 μm, Background Subtraction = true, “Quality” above 4000. These parameters were determined by trial and error during preliminary experiments. The Intensity Sum was found for each nucleus and the mean Intensity Sum was calculated for each aBLA. This entire experiment was replicated, with 19 mice in the first round and 23 mice in the second round. As mentioned previously, when the data were plotted, 1 sample in the second round had a mean Intensity Sum more than 2.5 SD greater than the mean for all 42 samples; this 1 poorly perfused sample was excluded from the analysis. For each round of the replication, the double-blind was broken only after the data analysis was complete.

For the male-female comparison in Figure 7C, the protocol was the same as above except there was no stress intervention.

#### Pharmacokinetics (Figure 6A)

##### LC-MS/MS Analysis

###### Sample preparation

Aliquots of 10 μL of plasma or 100 μL brain tissue homogenate (1 part tissue (g) + 2 parts water (mL)) were mixed with 10 μL of 100-nM Q-DEVD-OPh (internal standard) and 500 μL of deionized water. After liquid-liquid extraction (LLE) of analytes by 1 mL of dichloromethane, the lower layer was removed and 800 μL evaporated under a gentle stream of nitrogen. The residue was reconstituted in 100 μL of 70% mobile phase A/30% acetonitrile (see below); 5 μL (plasma) or 50 μL (brain homogenate) was injected into LC-MS/MS system.

###### LC conditions

Shimadzu 20A series liquid chromatography system; column: Agilent, Eclipse Plus C8, 3 x 50 mm, 1.8 μm particle size, at 45 C; mobile phase A: 98:2 H_2_O:acetonitrile (0.1% formic acid); mobile phase B: acetonitrile; elution gradient: 0-1 min 0-95% B, 1-1.5 min 95% B, 1.5-1.7 min 95-0% B; run time: 4 min.

###### ESI-MS/MS conditions

AB/SCIEX API 4000 QTrap tandem-mass spectrometer; positive mode electrospray (e.g., declustering potential, temperature, gas flow) and mass spectrometer (parent/daughter ions, quadrupole potential) parameters were optimized by direct infusion of pure analytes into the ESI-LC/MS system. MRM transitions (m/z units) monitored for quantification: Q-VD-OPh (Cayman Chemical, Cat#15260) 514.0/226.6 and Q-DEVD-OPh (internal standard) 800.3/256.5. Calibration samples in 0.38 – 1000 nM (plasma) and 0.5 – 61.3 nM (brain tissue) ranges were prepared by adding known amounts of pure Q-VD-OPh into plasma or brain tissue homogenate and were analyzed along with study samples. Linear response was observed in the concentration ranges measured (r=0.9997 and 0.9984) and the low limits of quantification (LLOQ) were 0.38 nM and 0.5 nM (+/-80% accuracy) in plasma and brain tissue, respectively.

#### Binding (Figure 6B)

NIMH’s Psychoactive Drug Screening Program (Besnard et al., 2012) determined the binding affinities of the receptors and transporters listed in Figure 6B.

#### Neurophysiology (Figure 6C-I)

##### Chronic Multi-Circuit Electrode Implants

All mice used in this study were 2-4-month-old male C57BL/6J from Jackson Labs. Mice were group housed (3-5 animals per cage) in a climate-controlled facility for rodents maintained on a 12-hour light-dark cycle and fed with food and water ad libitum. Electrodes were arranged in bundles and implanted as previously described (Dzirasa et al., 2013). Briefly, mice were anesthetized using isoflurane and placed in a stereotaxic frame. The isoflurane/oxygen flow rate was maintained at >1.5% saturation level and adjusted as needed to ensure full anesthesia as determined by tail-pinch response during the entire duration of surgery. Ground screws were placed in the skull above the cerebellum and the anterior cranium. HML coated tungsten wires were arranged in bundles and micro-wires were implanted into mice. All electrodes were referenced to the ground wire connected to the ground screws in the cranium. Electrode bundles were implanted as follows (all coordinates are measured from bregma):

| Brain area | AP coordinates (mm) | ML coordinates (mm) | DV coordinates (mm) |
| --- | --- | --- | --- |
| PrL | +1.8 | ±0.25 | −1.75 |
| Amyg | −1.46 | ±2.85 | −3.9 |
| DHipp | −2.4 | ±1.63 | −1.25 |

##### Neurophysiological Recordings

###### Acute drug treatment and brain network function

After 2-3 weeks of recovery from surgery, all mice were subjected to neurophysiological recordings. For 7 days, all mice were habituated to the handling required for connecting cables and performing saline injections. During the habituation period all mice were taken to the behavior room in their home cage and manually connected to the recording cables; after 15 minutes of recovery from handling, mice were given 20 minutes in the cage followed by a saline injection and another 60 minutes in the cage. After 7 days of habituation all mice were connected to the recording cables for the multi-channel Cerebus acquisition system (Blackrock Microsystems Inc., UT). Following 15 minutes of recovery from handling while connecting the recording cables, brain activities as local field potential (LFPs) (band-pass filtered at 0.3-250 Hz) were recorded and stored at 1000 Hz. LFP activity was recorded for 20 minutes while mice were in the cage before the saline/drug/DMSO injections. Behavioral data with automated tracking was also acquired (Neuromotive, Blackrock Microsystems Inc., UT). Data were analyzed using custom-written MATLAB scripts.

###### Brain network function before, during and after acute drug treatment

Following 7 days of habituation, all mice were subjected to neurophysiological recordings for 3 days as follows: first day, all mice were injected with saline; second day, 8 and 7 mice were injected with drug or DMSO, respectively; third day, all mice were injected with saline. All mice were recorded in their cage for 20 minutes prior to injections. Data were collected and stored for further analysis.

#### Statistical Analysis

For all mice, 20 minutes of home cage brain activity was compared with 20 minutes of brain activity following saline or drug/DMSO treatment. Four LFP channels for each structure were pseudorandomly chosen for analysis. LFP cross-structural coherence was calculated from the LFP pairs used for LFP oscillatory power analysis with the MATLAB (MathWorks, Natick, MA) mscohere function at a 1-second sliding window with a 1-second step.

LFPs and coherence activity were averaged for mice in the 6 frequencies that demonstrated bands of increased connectivity for 20 minutes: delta 1-4 Hz (Amy-Hip, Amy-PrL, and Hip-PrL), theta 4-11 Hz (Amy-Hip and Hip-PrL), and beta 15-30 Hz (PrL-Hip). The LFP power and coherence activity data were averaged across the recording session. Data were compared before and after treatment in the two groups using a RM-ANOVA with Box-Cox transformation, followed by a Wilcoxon sign-rank test with false discovery rate correction.

### QUANTIFICATION AND STATISTICAL ANALYSIS

Data analysis was performed with Microsoft Excel, Imaris (Bitplane), and MATLAB; and statistical analysis was done with GraphPad Prism 7. All statistical details can be found in the figure legends and figures. p<0.05 was the threshold of significance. Due to a poor perfusion, one data point in the experiment shown in Figure 7D was more than 2.5 SD greater than the mean for all 42 samples and was excluded from the analysis prior to unblinding.

## Results

We designed a bimolecular fluorescence complementation reporter of caspase activity that comprises four elements intended to maximize the signal-to-noise ratio: CAG promoters, split inteins, PEST sequences, and localization signals (Figure 1A, B). Bidirectional CAG promoters provide strong, ubiquitous expression (Niwa et al., 1991; Okabe et al., 1997) of each half of the mCerulean reporter (Rizzo et al., 2004). Split inteins further boost the signal by enabling covalent reconstitution of mCerulean from the two nonfluorescent halves (Wang et al., 2012). To minimize background signal, we designed these halves to be kept separate and continually degraded. This baseline state holds until active caspases cleave off the localization signals (Güttler et al., 2010) and PEST sequences (Li et al., 1998) by cutting canonical caspase cleavage sites (McStay et al., 2008) present between the mCerulean, localization, and PEST sequences. The liberated, stabilized mCerulean halves can then join via inteins and fluoresce, typically near the nucleus (Figure 1C). This reporter construct was introduced into the C57BL/6 mouse genome by pronuclear microinjection, yielding at least 24 copies in homozygotes according to qPCR. This line was bred onto a mixed background to increase the size and frequency of litters. After more than ten years of breeding, the copy number remained stable.

Apoptosis produces a strong signal in vivo and in vitro. Although the reporter is expected to produce background signal in mitotic cells, since only the PEST sequences would reduce nonspecific signal when the nuclear envelope disassembles, there is nevertheless substantial and significant separation of fluorescent signal between apoptotic and nonapoptotic cultured cells transiently transfected with the Apo plasmid (Figure 2A, B). As for postmitotic cells, application of 100-nM staurosporine to primary hippocampal cultures from transgenic mice caused a robust increase in signal that preceded cell death (Figure 3A). We caused in vivo apoptosis by injecting 6-hydroxydopamine into the medial forebrain bundle (He et al., 2000) and observed bright reporter signal and TUNEL-positive cells in the substantia nigra pars compacta (Figure 3B). We also induced apoptosis *in vivo* with middle cerebral artery occlusion (Beraldo et al., 2013) and found heightened cortical signal ipsilateral to the lesion (Figure 3C). In addition, we verified that a portion of reporter signal represents nonapoptotic caspase activity (NACA) by blocking its increase in acute hippocampal slices with a caspase inhibitor (Figure 3D; Caserta et al., 2003; Yang et al., 2004). Incidentally, in the hippocampus, NACA was mostly limited to neurons in the pyramidal layer (Figure 3E).

Due to the extreme brightness of reporter signal during apoptosis, we could measure it in the red portion of the spectrum (570-640 nm) even though mCerulean emits primarily in the cyan range (470-550 nm) (Table 1; Rizzo et al., 2004). Thus, it is simple to separate apoptotic (570-640 nm) from nonapoptotic or background (470-550 nm only) signal (see Methods: Apoptotic vs. Nonapoptotic Signal Intensity, Figure 4B). We confirmed that the prevalent baseline signal (Figure 4A, top row) was nonapoptotic since TUNEL stain was negative for the brightest cells (Figure 4C), despite being positive for shrunken epithelial cells in choroid plexus (Figure 4D). Likewise, bright signal was observed in the cerebellum of one-month-old *Arpc3* knockout mice, which demonstrate ataxia and loss of Purkinje cells at this age (Figure 4E; Kim et al., 2013). Of note, reporter signal is evident in shedding skin and between the toes of P0 pups (Figure 4F).

We further characterized the nonapoptotic signal, or NACA, in whole-brain scans using different imaging modalities (see Table S1). NACA was widespread and especially prominent in the amygdala, hippocampus, cortex, and Purkinje cells (Figure 5A-C). Importantly, *in situ* hybridization showed a similar pattern of mCerulean expression (Figure 5D), suggesting that a portion of baseline NACA is due to nonspecific background reporter signal. Therefore, bringing the above findings together, we have generated a transgenic reporter of caspase activity that can differentiate apoptotic from nonapoptotic activity for neuronal mapping studies throughout the brain.

To determine which brain regions may have functionally relevant NACA, we measured the effect of a caspase inhibitor on *in vivo* neurophysiology. We first determined that the inhibitor Q-VD-OPh reaches IC50 levels (Caserta et al., 2003) in brain when administered intraperitoneally at 5 mg/kg (Figure 6A) and shows no off-target activity at potential psychoactive targets at these concentrations (Figure 6B). Comparing multielectrode recordings of local field potentials (Dzirasa et al., 2013) in the basal amygdala, dorsal hippocampus, and prelimbic cortex 24 hours before treatment and 24 hours after treatment (Figure 6C-E), we detected a reduction in 15-30-Hz functional connectivity between the amygdala and prelimbic cortex (Figure 6F, G) and a specific reduction in 15-30-Hz amplitude in the amygdala (Figure 6H, I). Hence, NACA may play an important role in neuronal function in the aBLA.

Because of the importance of the aBLA in behavioral stress (Roozendaal et al., 2009; Kim et al., 2016, 2017), we tested whether NACA was related to stress responses in the amygdala. We first attempted to determine the cell type responsible for the majority of nonapoptotic reporter signal in the amygdala. Our immunohistological assays suggest that neurons – likely excitatory principal neurons – are the primary source of NACA-related signal in the aBLA (Figure 7A, B; Neely et al., 2009). Next, we noted that there was a baseline difference in NACA levels between males and females (Figure 7C) and therefore studied both sexes concurrently in a recently reported stress paradigm (Chu et al., 2016). Nine days after a 24-hour bout of restraint stress, we unexpectedly found a persistent increase in aBLA NACA only in females (Figure 7D). Therefore, NACA may be a key component of stress responsiveness in the amygdala, especially in females.

## Discussion

We designed a transgenic fluorescent reporter sensitive and specific enough to detect NACA as well as caspase-mediated apoptosis in mouse brain. Due to the logarithmic increase in caspase activity during apoptosis (Li et al., 2010), it was possible to separate long-wavelength apoptotic signal from short-wavelength nonapoptotic signal. We obtained evidence of a causal relationship between baseline NACA and neural activity by measuring functional connectivity in unstressed mice treated with a caspase inhibitor. Caspase inhibition caused a reduction in functional connectivity between the basal amygdala and prelimbic cortex, along with reduced 15-30-Hz power in the amygdala. Finally, we characterized NACA in the aBLA by relating it to stress responsiveness. We uncovered a sex-specific effect of stress on NACA: 9 days after prolonged restraint stress, NACA was elevated only in female mice. Together these results demonstrate the utility of our caspase reporter mouse and the potential importance of NACA in studies of neural function and stress.

Our primary aim was to design a transgenic reporter that would enable mapping of all caspase activity across the brain using multiphoton and other forms of microscopy. Consequently, our reporter contains several design elements subserving this goal that in other contexts might constitute limitations or shortcomings. (1) There is no internal control, such as a contrasting, constitutively expressed fluorescent protein (Yamaguchi et al., 2011), because multiphoton absorption spectra typically overlap (Drobizhev et al., 2011), and any subsequent emission overlap from a bright control would impair the potentially weak signal of the reporter. (2) The reporter does not indicate the subcellular location of caspase activity, since, to facilitate mapping and cellwise quantification, we wanted peak signal to center on the nucleus regardless of the source of caspase activity. (3) Reporter signal is not specifically targeted to the nucleus because fluorescence in the soma can demonstrate shrinkage during apoptosis. (4) The cyan fluorophore overlaps the autofluorescence profile of neural tissue (Billinton & Knight, 2001). This design element enables crossing to other reporter strains, such as calcium sensor mice, which utilize green or red fluorescent proteins. In addition, the tail-end red emission of our mCerulean reporter during apoptosis can be detected by multiphoton microscopy, unlike the near-infrared tail-end emission of a red fluorophore, thereby facilitating separation of nonapoptotic from apoptotic signal. (5) The reconstituted reporter has a long half-life – probably >1 day (Corish & Tyler-Smith, 1999) – since a destabilized fluorescent protein would reduce our chances of detecting transient, weak caspase activity. (6) Our system is not controlled by Cre or other recombinases because this would limit genomic copy number to one, or perhaps two, which would constrain reporter signal output and make detection of weak caspase activity less likely. (7) Likewise, the insertion site of our reporter construct is unknown, since we generated the mouse by pronuclear microinjection. But this random insertion enabled us to reach a homozygous copy number of at least 24, according to quantitative PCR, which has been stable for over ten years of breeding. (8) Lastly, with such a sensitive reporter of caspase activity, tissue fixation must be performed with great care to maintain consistency of NACA signal and to avoid artifacts like “dark neurons” (Beevor, 1883; Cammermeyer, 1960; Jortner, 2006). In order to obtain reproducible results, we strongly recommend following the fixation procedure described in the Methods.

Aside from the various limitations by design, the caspase activity reporter has a few important caveats. First, the baseline fluorescence of the transgenic reporter is nonzero; that is, healthy wild-type adult brains always contain some faint fluorescent reporter signal in some regions and cell types. Our data suggest that this baseline signal is a mixture of specific, caspase-induced fluorescence and nonspecific, caspase-independent fluorescence, the latter most likely due to robust activity of the CAG promoter in certain brain regions and cell types. Therefore, it is incumbent on the researcher to include a control group in every experiment to account for the unknown nonspecific reporter signal in these transgenic mice. It is important to note, however, that in over ten years of working with these mice, we have not detected bright, apoptosis-associated signal in healthy adult brain, except occasionally in the choroid plexus. Second, since one of the signal-quenching mechanisms of the reporter depends on an intact nuclear envelope, one would expect to see some fluorescent signal in dividing cells – although this type of signal would be limited by the rapid degradation imparted by intact PEST sequences. Due to this spurious fluorescence associated with cell division, the caspase reporter is best suited to the study of postmitotic tissues. Lastly, since we have not seen baseline apoptotic signal – or TUNEL-positive cells – in healthy adult brain in our decade of research on these mice, we doubt that the transgenic reporter has a toxic effect on cells. Indeed, the baseline fluorescent signal seen in brain is typically weak and difficult to distinguish from autofluorescence; hence, it is unlikely that the reporter is disturbing proteostasis, and we have seen no evidence of bright fluorescent clumps or aggregates. But we have not performed a rigorous battery of tests to demonstrate a lack of any cellular toxicity.

When we applied our careful sample preparation and microscopy techniques to the study of stress, we found a sex-specific persistent increase in NACA in the aBLA in females. Although surprising, if we suppose that caspases play a meaningful role in stress responsiveness, this result is not entirely unexpected. As mentioned above, the caspase pathway is engaged in apoptotic cell death in female brain, but a parallel cascade predominates in male brain (Zhu et al., 2006; Liu et al., 2009). Also, long-lived physiologic changes after stress tend to be confined to the amygdala. Perhaps the most widely reported alteration in the amygdala after stress is persistent dendritic hypertrophy (Roozendaal et al., 2009). Furthermore, neurons in the aBLA fire more after stress and drive negative-valence defensive behaviors (Kim et al., 2016, 2017). Hence, caspase activity may help explain alterations in neuronal morphology and activity, and, at least in females, the peculiar structural and functional responses to stress in the amygdala.

In conclusion, we have developed a transgenic reporter of caspase activity in mice, as well as tissue fixation and microscopy methods that enable the mapping and quantification of weak and transient nonapoptotic caspase activity. We obtained evidence of a causal relation between caspase activity and amygdalar functional connectivity; and we utilized our system to uncover a sex-dependent effect of stress on the anterior basolateral amygdala. Our reporter and techniques can be used to test hypotheses relating to apoptotic and nonapoptotic caspase activity throughout the brain and the rest of the organism.

## Author Contributions

PJN, TFP, NMU, SK, YZ, GI, JDG, GT, VFP, MAMP, IS, KD, GAJ, and MGC designed research. PJN, TFP, NMU, SK, YZ, GI, JDG, GT, PF, VGO, MSG, FB, JH, and IS performed research. PJN, TFP, JDG, WLR, IS, and JCS contributed new reagents. PJN, EC, JCS, and KD contributed analytic tools. PJN, SK, GI, JDG, PF, VGO, JH, IS, and KD analyzed data. PJN wrote the paper.

## Acknowledgments

We are grateful to the following for their assistance and advice: Rainbo Hultman, Benjamin Sachs, Lauren Rochelle, Daniel Urban, Scott Robertson, Jue Fan, Nicole Calakos, Fan Wang, David Piston, Martin Fischer, Francisco Robles Guerrero, Ramona Rodriguiz, William Wetsel, Cheryl Bock, Meilang Flowers, Thomas L. Brown, Lauren Burianek, and Scott Soderling. Ki determinations were provided by the NIMH’s Psychoactive Drug Screening Program, Contract #HHSN-271-2013-00017-C. Pharmacokinetic analysis was carried out by the Duke Cancer Institute PK/PD Core Laboratory. This work was supported by NIH grants: 1R21NS081513 (PJN), 8P41EB015897 (GAJ), 2R01MH079201 and 5R37MH073853 (MGC); and CIHR grants: MOP 93651, 12600, 89919 (MAMP). Support from the Pall Family Foundation and Lennon Family Foundation to MGC is greatly appreciated.

## Conflict of Interest

A. No, authors report no conflict of interest.

## Funding sources

This work was supported by NIH grants: 1R21NS081513 (PJN), 8P41EB015897 (GAJ), 2R01MH079201 and 5R37MH073853 (MGC); and CIHR grants: MOP 93651, 12600, 89919 (MAMP). Support from the Pall Family Foundation and Lennon Family Foundation to MGC is greatly appreciated.

## Notes

### Competing Interest Statement

The authors have declared no competing interest.

### Summary of Updates

Alterations in response to reviewer comments

## References

Bardet, P.L., Kolahgar, G., Mynett, A., Miguel-Aliaga, I., Briscoe, J., Meier, P., and Vincent, J.P. (2008). A fluorescent reporter of caspase activity for live imaging. Proc. Natl. Acad. Sci. U.S.A. 105, 13901–13905.

Beevor, C. (1883). Die Kleinhirnrinde. Arch. Amt. u. Physiol., An. & Abt. (no vol.), 365–388.

Beraldo, F.H., Arantes, C.P., Santos, T.G., Queiroz, N.G., Young, K., Rylett, R.J., Markus, R.P., Prado, M.A., and Martins, V.R. (2010). Role of alpha7 nicotinic acetylcholine receptor in calcium signaling induced by prion protein interaction with stress-inducible protein 1. J. Biol. Chem. 285, 36542–36550.

Beraldo, F.H., Soares, I.N., Goncalves, D.F., Fan, J., Thomas, A.A., Santos, T.G., Mohammad, A.H., Roffé, M., Calder, M.D., Nikolova, S., et al. (2013). Stress-inducible phosphoprotein 1 has unique cochaperone activity during development and regulates cellular response to ischemia via the prion protein. FASEB J. 27, 3594–3607.

Besnard, J., Ruda, G.F., Setola, V., Abecassis, K., Rodriguiz, R.M., Huang, X.P., Norval, S., Sassano, M.F., Shin, A.I., Webster, L.A., et al. (2012). Automated design of ligands to polypharmacological profiles. Nature 492, 215–220.

Billinton, N., and Knight, A.W. (2001). Seeing the wood through the trees: a review of techniques for distinguishing green fluorescent protein from endogenous autofluorescence. Anal Biochem. 291, 175–197.

Cammermeyer, J. (1960). The post-mortem origin and mechanism of neuronal hyperchromatosis and nuclear pyknosis. Exp. Neurol. 2, 379–405.

Campbell, D.S., and Okamoto, H. (2013). Local caspase activation interacts with Slit-Robo signaling to restrict axonal arborization. J. Cell Biol. 203, 657–672.

Caserta, T.M., Smith, A.N., Gultice, A.D., Reedy, M.A., and Brown, T.L. (2003). Q-VD-OPh, a broad spectrum caspase inhibitor with potent antiapoptotic properties. Apoptosis 8, 345–352.

Chen, S.X., Cherry, A., Tari, P.K., Podgorski, K., Kwong, Y.K., and Haas, K. (2012). The transcription factor MEF2 directs developmental visually driven functional and structural metaplasticity. Cell 151, 41–55.

Chu, X., Zhou, Y., Hu, Z., Lou, J., Song, W., Li, J., Liang, X., Chen, C., Wang, S., Yang, B., et al. (2016). 24-hour-restraint stress induces long-term depressive-like phenotypes in mice. Sci. Rep. 6, 32935.

Corish, P., and Tyler-Smith, C. (1999). Attenuation of green fluorescent protein half-life in mammalian cells. Protein Eng. 12, 1035–1040.

Drobizhev, M., Makarov, N.S., Tillo, S.E., Hughes, T.E., and Rebane, A. (2011). Two-photon absorption properties of fluorescent proteins. Nat. Methods 8, 393–399.

Du, L., Bayir, H., Lai, Y., Zhang, X., Kochanek, P.M., Watkins, S.C., Graham, S.H., and Clark, R.S. (2004). Innate gender-based proclivity in response to cytotoxicity and programmed cell death pathway. J. Biol. Chem. 279, 38563–38570.

Dzirasa, K., Kumar, S., Sachs, B.D., Caron, M.G., and Nicolelis, M.A. (2013). Cortical-amygdalar circuit dysfunction in a genetic mouse model of serotonin deficiency. J. Neurosci. 33, 4505–4513.

Ertürk, A., Wang, Y., and Sheng, M. (2014). Local pruning of dendrites and spines by caspase-3-dependent and proteasome-limited mechanisms. J. Neurosci. 34, 1672–1688.

Evans, T.C. Jr, Martin, D., Kolly, R., Panne, D., Sun, L., Ghosh, I., Chen, L., Benner, J., Liu, X.Q., and Xu, M.Q. (2000). Protein trans-splicing and cyclization by a naturally split intein from the dnaE gene of Synechocystis species PCC6803. J. Biol. Chem. 275, 9091–9094.

Fernando, P., Brunette, S., and Megeney, L.A. (2005). Neural stem cell differentiation is dependent upon endogenous caspase 3 activity. FASEB J. 19, 1671–1673.

Fu, Q., Duan, X., Yan, S., Wang, L., Zhou, Y., Jia, S., Du, J., Wang, X., Zhang, Y., and Zhan, L. (2013). Bioluminescence imaging of caspase-3 activity in mouse liver. Apoptosis 18, 998–1007.

Galbán, S., Jeon, Y.H., Bowman, B.M., Stevenson, J., Sebolt, K.A., Sharkey, L.M., Lafferty, M., Hoff, B.A., Butler, B.L., Wigdal, S.S., et al. (2013). Imaging proteolytic activity in live cells and animal models. PLoS One 8, e66248.

Gogarten, J.P., Senejani, A.G., Zhaxybayeva, O., Olendzenski, L., and Hilario, E. (2002). Inteins: structure, function, and evolution. Annu. Rev. Microbiol. 56, 263–287.

Güttler, T., Madl, T., Neumann, P., Deichsel, D., Corsini, L., Monecke, T., Ficner, R., Sattler, M., and Görlich, D. (2010). NES consensus redefined by structures of PKI-type and Rev-type nuclear export signals bound to CRM1. Nat. Struct. Mol. Biol. 17, 1367–1376.

Hama, H., Kurokawa, H., Kawano, H., Ando, R., Shimogori, T., Noda, H., Fukami, K., Sakaue-Sawano, A., and Miyawaki, A. (2011). Scale: a chemical approach for fluorescence imaging and reconstruction of transparent mouse brain. Nat. Neurosci. 14, 1481–1488.

He, Y., Lee, T., and Leong, S.K. (2000). 6-Hydroxydopamine induced apoptosis of dopaminergic cells in the rat substantia nigra. Brain Res. 858, 163–166.

Jiao, S., and Li, Z. (2011). Nonapoptotic function of BAD and BAX in long-term depression of synaptic transmission. Neuron 70, 758–772.

Jortner, B.S. (2006). The return of the dark neuron. A histological artifact complicating contemporary neurotoxicologic evaluation. Neurotoxicology 27, 628–634.

Khanna, D., Hamilton, C.A., Bhojani, M.S., Lee, K.C., Dlugosz, A., Ross, B.D., and Rehemtulla, A. (2010). A transgenic mouse for imaging caspase-dependent apoptosis within the skin. J. Invest. Dermatol. 130, 1797–1806.

Kim, I.H., Racz, B., Wang, H., Burianek, L., Weinberg, R., Yasuda, R., Wetsel, W.C., and Soderling, S.H. (2013). Disruption of Arp2/3 results in asymmetric structural plasticity of dendritic spines and progressive synaptic and behavioral abnormalities. J. Neurosci. 33, 6081–6092.

Kim, J., Pignatelli, M., Xu, S., Itohara, S., and Tonegawa, S. (2016). Antagonistic negative and positive neurons of the basolateral amygdala. Nat. Neurosci. 19, 1636–1646.

Kim, J., Zhang, X., Muralidhar, S., LeBlanc, S.A., and Tonegawa, S. (2017). Basolateral to Central Amygdala Neural Circuits for Appetitive Behaviors. Neuron 93, 1464–1479.e5.

Lennon, C.W., and Belfort, M. (2017). Inteins. Curr. Biol. 27, R204–R206.

Li, X., Zhao, X., Fang, Y., Jiang, X., Duong, T., Fan, C., Huang, C.C., and Kain, S.R. (1998). Generation of destabilized green fluorescent protein as a transcription reporter. J. Biol. Chem. 273, 34970–34975.

Li, Z., Jo, J., Jia, J.M., Lo, S.C., Whitcomb, D.J., Jiao, S., Cho, K., and Sheng, M. (2010). Caspase-3 activation via mitochondria is required for long-term depression and AMPA receptor internalization. Cell 141, 859–871.

Liu, F., Li, Z., Li, J., Siegel, C., Yuan, R., and McCullough, L.D. (2009). Sex differences in caspase activation after stroke. Stroke 40, 1842–1848.

Liu, F., Lang, J., Li, J., Benashski, S.E., Siegel, M., Xu, Y., and McCullough, L.D. (2011). Sex differences in the response to poly(ADP-ribose) polymerase-1 deletion and caspase inhibition after stroke. Stroke 42, 1090–1096.

McStay, G.P., Salvesen, G.S., and Green, D.R. (2008). Overlapping cleavage motif selectivity of caspases: implications for analysis of apoptotic pathways. Cell Death Differ. 15, 322–331.

Neely, M.D., Robert, E.M., Baucum, A.J., Colbran, R.J., Muly, E.C., and Deutch, A.Y. (2009). Localization of myocyte enhancer factor 2 in the rodent forebrain: regionally-specific cytoplasmic expression of MEF2A. Brain Res. 1274, 55–65.

Niwa, H., Yamamura, K., and Miyazaki, J. (1991). Efficient selection for high-expression transfectants with a novel eukaryotic vector. Gene 108, 193–199.

Okabe, M., Ikawa, M., Kominami, K., Nakanishi, T., and Nishimune, Y. (1997). ’Green mice’ as a source of ubiquitous green cells. FEBS Lett. 407, 313–319.

Renolleau, S., Fau, S., Goyenvalle, C., Joly, L.M., Chauvier, D., Jacotot, E., Mariani, J., and Charriaut-Marlangue, C. (2007). Specific caspase inhibitor Q-VD-OPh prevents neonatal stroke in P7 rat: a role for gender. J. Neurochem. 100, 1062–1071.

Rizzo, M.A., Springer, G.H., Granada, B., and Piston, D.W. (2004). An improved cyan fluorescent protein variant useful for FRET. Nat. Biotechnol. 22, 445–449.

Roozendaal, B., McEwen, B.S., and Chattarji, S. (2009). Stress, memory and the amygdala. Nat. Rev. Neurosci. 10, 423–433.

Shah, N.H., and Muir, T.W. (2014). Inteins: Nature’s Gift to Protein Chemists. Chem. Sci. 5, 446–461.

Tang, H.L., Tang, H.M., Fung, M.C., and Hardwick, J.M. (2015). In vivo CaspaseTracker biosensor system for detecting anastasis and non-apoptotic caspase activity. Sci. Rep. 5, 9015.

Unsain, N., and Barker, P.A. (2015). New Views on the Misconstrued: Executioner Caspases and Their Diverse Non-apoptotic Roles. Neuron 88, 461–474.

Wang, P., Chen, T., Sakurai, K., Han, B.X., He, Z., Feng, G., and Wang, F. (2012). Intersectional Cre driver lines generated using split-intein mediated split-Cre reconstitution. Sci. Rep. 2, 497.

Westphal, D., Sytnyk, V., Schachner, M., and Leshchyns’ka, I. (2010). Clustering of the neural cell adhesion molecule (NCAM) at the neuronal cell surface induces caspase-8- and -3-dependent changes of the spectrin meshwork required for NCAM-mediated neurite outgrowth. J. Biol. Chem. 285, 42046–42057.

Wu, H., Hu, Z., and Liu, X.Q. (1998). Protein trans-splicing by a split intein encoded in a split DnaE gene of Synechocystis sp. PCC6803. Proc. Natl. Acad. Sci. USA. 95, 9226–9231.

Yamaguchi, Y., Shinotsuka, N., Nonomura, K., Takemoto, K., Kuida, K., Yosida, H., and Miura, M. (2011). Live imaging of apoptosis in a novel transgenic mouse highlights its role in neural tube closure. J. Cell Biol. 195, 1047–1060.

Yang, L., Sugama, S., Mischak, R.P., Kiaei, M., Bizat, N., Brouillet, E., Joh, T.H., and Beal, M.F. (2004). A novel systemically active caspase inhibitor attenuates the toxicities of MPTP, malonate, and 3NP in vivo. Neurobiol. Dis. 17, 250–259.

Yuan, J., Shaham, S., Ledoux, S., Ellis, H.M., and Horvitz H.R. (1993). The C. elegans cell death gene ced-3 encodes a protein similar to mammalian interleukin-1 beta-converting enzyme. Cell 75, 641–652.

Zhu, C., Xu, F., Wang, X., Shibata, M., Uchiyama, Y., Blomgren, K., and Hagberg, H. (2006). Different apoptotic mechanisms are activated in male and female brains after neonatal hypoxia-ischaemia. J. Neurochem. 96, 1016–1027.

